# Deactivation of ligand-receptor interactions enhancing lymphocyte infiltration drives melanoma resistance to Immune Checkpoint Blockade

**DOI:** 10.1101/2023.09.20.558683

**Authors:** Sahil Sahni, Binbin Wang, Di Wu, Saugato Rahman Dhruba, Matthew Nagy, Sushant Patkar, Ingrid Ferreira, Kun Wang, Eytan Ruppin

## Abstract

Immune checkpoint blockade (ICB) is a promising cancer therapy; however, resistance often develops. To learn more about ICB resistance mechanisms, we developed IRIS (**I**mmunotherapy **R**esistance cell-cell **I**nteraction **S**canner), a machine learning model aimed at identifying candidate ligand-receptor interactions (LRI) that are likely to mediate ICB resistance in the tumor microenvironment (TME). We developed and applied IRIS to identify resistance-mediating cell-type-specific ligand-receptor interactions by analyzing deconvolved transcriptomics data of the five largest melanoma ICB therapy cohorts. This analysis identifies a set of specific ligand-receptor pairs that are deactivated as tumors develop resistance, which we refer to as *resistance deactivated interactions (RDI).* Quite strikingly, the activity of these RDIs in pre-treatment samples offers a markedly stronger predictive signal for ICB therapy response compared to those that are activated as tumors develop resistance. Their predictive accuracy surpasses the state-of-the-art published transcriptomics biomarker signatures across an array of melanoma ICB datasets. Many of these RDIs are involved in chemokine signaling. Indeed, we further validate on an independent large melanoma patient cohort that their activity is associated with CD8+ T cell infiltration and enriched in hot/brisk tumors. Taken together, this study presents a new strongly predictive ICB response biomarker signature, showing that following ICB treatment resistant tumors turn inhibit lymphocyte infiltration by deactivating specific key ligand-receptor interactions.

## Introduction

Immune checkpoint blockade (ICB) therapy provides durable clinical benefits to melanoma patients, but the response rate remains limited, with approximately 40% of patients experiencing positive outcomes (1). The majority of patients receiving checkpoint therapy, both in the case of primary treatment or subsequent treatment after initial response, face the challenge of developing resistance (2,3). The mechanisms of ICB resistance are complex and still not well understood. Current hypothesized mechanisms of resistance include depletion of neoantigen expression by tumor cells, deficiencies in antigen-presentation machinery, and premature exhaustion of effector T cells (4). Moreover, the composition of the tumor microenvironment (TME) can alter ICB response by composing an immunosuppressive or immunoreactive TME (4–6). Identifying resistance mechanisms is crucial for developing combination therapies that target multiple resistance mediators simultaneously, improving patient response to ICB (4).

Numerous transcriptomic-based biomarkers have emerged for predicting ICB therapy response and uncovering mechanisms of resistance. *TIDE* infers gene signatures associated with T cell dysfunction and exclusion from TCGA using bulk transcriptomics (7). *IMPRES* learned the pairwise relations of 15 checkpoint genes relevant to spontaneous regression in neuroblastoma using bulk transcriptomics (8). *MPS* uncovered a melanocytic plasticity signature associated with ICB therapy resistance, derived from a combination of mouse models and bulk transcriptomics of ICB patients (9). *Davoli et al.* identified a gene-expression signature correlated with aneuploidy and negative correlated with immune infiltration for melanoma patients using bulk transcriptomics (10). *Jerby-Arnon et al.* elucidated a transcriptomic program associated with T cell exclusion and immune evasion from scRNA-seq of ICB-treated melanoma patients (11). Tres identifies signatures of T cells that are resilient to immunosuppressive TME and confer antitumor properties from single-cell transcriptomics (12). However, these biomarkers have notable limitations: 1. Bulk transcriptomic biomarkers overlook cell type heterogeneity within TME; 2. Single-cell transcriptomic signatures may not apply to the much more abundant bulk ICB cohorts; 3. Most of the pertaining genes composing these biomarker signatures are not targetable; and 4. Their predictive power leaves room for further improvement.

Cell-cell interactions in the TME plays a crucial role in impacting tumor growth and clinical outcome (13). As a fundamental signaling mechanism, it is mediated by cell-type-specific ligand-receptor interactions, which ultimately underlie ICB therapy (4,5,13). Characterization of ICB therapy resistance mediating interactions is likely to advance our understanding of the development of resistance, enhance ICB response prediction, and help discover novel therapeutic targets. However, established approaches for profiling cell-type-specific gene expression (e.g., FACS) do not directly provide such interaction data (14). To dissect bulk transcriptomes and prioritize clinically relevant cell-cell interactions, we recently developed CODEFACS (COnfident DEconvolution for All Cell Subsets) and LIRICS (LIgand-Receptor Interactions between Cell Subsets), respectively (14). These methods have laid the basis for the development here of IRIS (**I**mmunotherapy **R**esistance cell-cell **I**nteraction **S**canner), a supervised machine learning method for identifying de-novo ligand-receptor interactions relevant to ICB therapy response in the TME.

We applied IRIS to identify resistance-mediating cell-type-specific ligand-receptor interactions in the five largest ICB therapy cohorts in melanoma (8,15–18). Through this analysis, we discovered that specific ligand-receptor pairs are deactivated as tumors develop resistance, which we refer to as *resistance deactivated interactions (RDI).* The activity state of these RDIs is predictive of treatment response across melanoma ICB datasets, surpassing previously published transcriptomics biomarkers. Many RDIs are involved in chemokine signaling, primarily associated with the recruitment of CD8+ T cells to the TME. Their activity is associated with CD8+ T cell infiltration and enriched in hot (brisk) tumors compared to cold (non-brisk) ones. These findings suggest that following ICB treatment, resistant tumors turn their TMEs cold through the inhibition of lymphocyte infiltration by deactivating specific ligand-receptor interactions.

## Results

### Overview of IRIS and the analysis

We obtained three deconvolved melanoma ICB cohorts (15–17) from the CODEFACS publication and further deconvolved two additional melanoma ICB cohorts (8,18) using CODEFACS, a deconvolution method recently developed by our laboratory (14). The expression profile of ten cell types in the TME (B cells, CD8+ T cells, CD4+ T cells, cancer associated fibroblasts, endothelial cells, macrophages, malignant cells, NK cells, plasmacytoid dendritic cells, and skin dendritic cells) are derived for each patients’ tumor sample from the ICB cohorts. Subsequently, we employed LIRICS (14), a ligand-receptor interaction inference tool, to derive cell-type-specific ligand-receptor interaction activity profile in each patient given the output from CODEFACS. The corresponding clinical information including response labels, survival and timelines were available for each patient.

We developed IRIS (**I**mmunotherapy **R**esistance cell-cell **I**nteraction **S**canner), a computational method specifically designed to identify immune checkpoint blockade (ICB) resistance relevant ligand-receptor interactions in the tumor microenvironment (TME), given a patients cohort including tumor bulk expression data and ICB treatment response data. The gene expression data is deconvolved using CODEFACS such that the input to IRIS in a given patients cohort is comprised of two components (**Fig. 1**): 1. cell-type-specific ligand-receptors interaction activity profiles (denoting either activation: 1 or inactivation: 0) in each tumor sample, which is inferred using LIRICS from the deconvolved expression – an interaction is considered as activated if the (deconvolved) expression of both its ligand and receptor genes is above their median expression values across the cohort samples, and deactivated otherwise; 2. The corresponding ICB response outcome for each patient.

**Figure 1:**
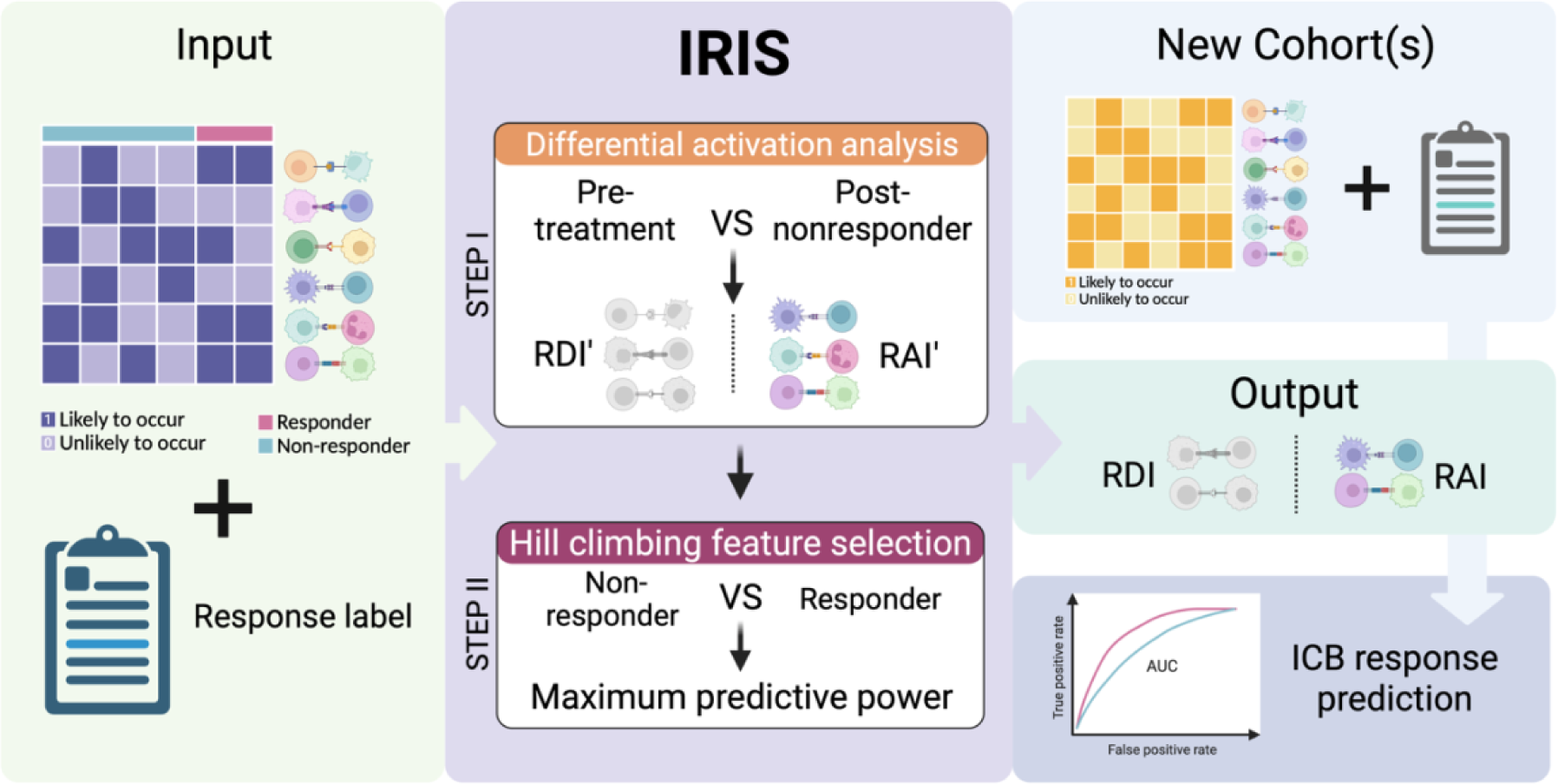
Overview of IRIS. The IRIS input includes cell-type-specific ligand-receptor activity profiles (inferred via applying CODEFACS and LIRICS on the tumor transcriptomics) and treatment response labels. It consists of two steps: Step I use a Fisher’s test to identify differentially activated ligand-receptor interactions in pre-treatment and non-responder post-treatment samples. These interactions are categorized as either resistant deactivated interactions (RDI) or resistant activated interactions (RAI) based on their differential activity state in the post-treatment vs. the pre-treatment state; that is, RDIs are deactivated in post-treatment resistant patients and vice versa for RAIs. Step II employs a hill climbing aggregative feature selection algorithm to choose the optimal set of RDIs or RAIs for classifying responders and non-responders in pre-treatment samples. The final output of IRIS is a selected set of RDIs and RAIs hypothesized to facilitate in ICB resistance, that can be used to predict ICB therapy response in a new ICB cohort.

Based on both the ligand-receptor interaction activity and response data, IRIS employs a two-step supervised machine learning (ML) method to identify ICB resistance relevant interactions (**Fig. 1**). (***1***) In the first step, the model selects interactions that exhibit differential activation between pre-treatment and on-treatment *non-responder* (progressive/stable disease) patients. The primary objective of this step is to identify interactions whose activation state following exposure to ICB is associated with the development of resistance. These interactions are categorized as *resistant deactivated interactions (RDI)* or *resistant activated interactions (RAI),* based on their differential activity state in the post-treatment vs. the pre-treatment samples; that is, RDIs are deactivated in post-treatment resistant patients and vice versa for RAIs. (***2***) In the second step, we employ a hill climbing aggregative feature selection algorithm to select an optimal set of LRIs that maximizes the classification power in distinguishing responders and non-responders from the pre-treatment tumor transcriptomics. When multiple training cohorts are available, both steps are executed iteratively on each cohort, resulting in cohort-specific models – an optimal set of response predictive LRIs for each cohort, which are composed of resistance deactivated interactions (RDI) and resistance activated ones (RAI). Additionally, each patient’s *resistant activated score (RAS)* and *resistant deactivated score (RDS)* is computed for each tumor sample. RAS represents the normalized count of active RAIs, while RDS represents the normalized count of active RDIs. Higher RAS indicates non-responsiveness, while higher RDS indicates higher responsiveness to ICB therapy – the more active-state RDIs present in a tumor in the pre-treatment stage, the more likely is the patient to respond to the ICB therapy.

### Deactivated interactions in resistant patients offer markedly stronger predictive value for ICB therapy response compared to activated ones

Functional resistance relevant interactions are anticipated to be predictive of ICB therapy response. To evaluate and quantify their response prediction power, we employed a “leave one cohort out” strategy to assess the performance of RAS and RDS scores in predicting ICB response. This involves iteratively designating each of the five cohorts as the *independent test cohort* while utilizing the remaining ones as the training cohorts to identify a set of resistance relevant interactions using IRIS. Furthermore, the identified RAIs and RDIs from all the training cohorts, in aggregate, are utilized to derive RAS and RDS scores for ICB response prediction in the left-out test cohort.

Strikingly, we observe that the RDS significantly outperforms RAS in predicting ICB therapy response (Wilcoxon 1-sided paired test p-value: 0.0039, **Fig. 2A**). The mean area under the curve (AUC) over all 5 independent test cohort for RDS is 0.72, while for RAS it is a dismal 0.39 (**Supp Fig. 1A**). Furthermore, we conducted a more in-depth assessment on the association between each individual interaction and ICB response in pre-treatment samples and post-treatment samples separately. The results reveal that RDIs are significantly more enriched for interactions involved in immune processes compared to RAIs across both pre-treatment and post-treatment samples (Fisher’s exact test: p-value < 2.2e-16 and < 2.2e-16; odds ratio: ∼18.8 and ∼31.1 respectively) (**Supp Fig. 4B-E**). These findings highlight the potential functional importance of RDIs in mediating ICB resistance, in contrast to the RAIs that lack predictive power. Therefore, we focus on RDIs in our subsequent analyses.

**Figure 2:**
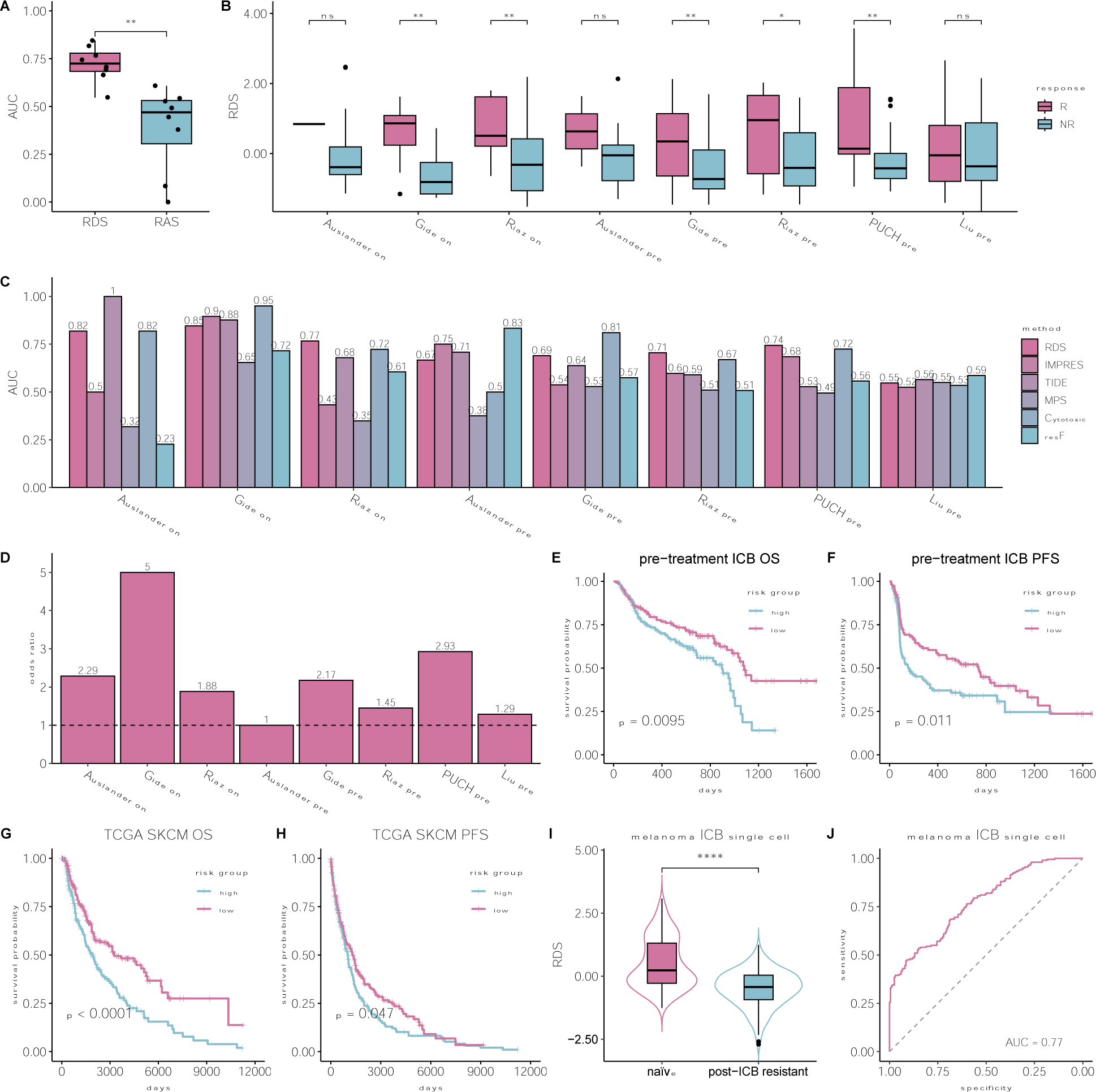
RDS based prediction of ICB response in melanoma. **(A)** Boxplot depicting the distribution of AUCs in classifying responder vs. non-responder samples in all melanoma ICB cohorts between inferred resistant deactivated (RDI) or resistant activated interactions (RAI). **(B)** Boxplot depicting the distribution of resistance deactivated score (RDS) between responder (R) and non-responder (NR) melanoma ICB samples. **(C)** Bar plot depicting the AUC in classifying responder vs. non-responder melanoma samples for numerous published transcriptomic based prediction scores including RDS, IMPRES (8), TIDE (7), MPS (9), Cytotoxic signature (10), and functional ICB resistance (resF) (11). **(D)** Bar plots depicting the odds-ratio in classifying true responder and non-responder samples using RDS scores. **(E)** Kaplan-Meier plot depicting overall survival of the combined set of pre-treatment melanoma samples receiving immune checkpoint blockade (N=274). The patients are stratified into low-risk/high-risk groups based on the RDS median value. **(F)** Kaplan-Meier plot depicting progression-free survival of the combined set of pre-treatment melanoma samples receiving immune checkpoint blockade (N=206). The patients are stratified into low-risk/high-risk groups as above. **(G)** Kaplan-Meier plot depicting overall survival of the combined set of pre-treatment TCGA-SKCM samples (N=438). The patients are stratified into low-risk/high-risk groups as above. **(H)** Kaplan-Meier plot depicting progression-free survival of the combined set of pre-treatment TCGA-SKCM samples (N=423). The patients are stratified into low-risk/high-risk groups as above. **(I)** Boxplot depicting the distribution of RDS between untreated (naïve) and post-checkpoint resistance (post-ICB resistant) samples from a melanoma ICB single-cell cohort. **(J)** ROC curve depicting the classification accuracy of naïve vs. post-ICB resistant tumors in a melanoma ICB single-cell cohort based on the RDS scores of each patient sample (**Methods**).

Further assessing the predictive power of RDS we find that responders exhibit significantly higher RDS compared to non-responders across five cohorts (**Fig. 2B**). Additionally, we find that the predictive power of RDS (average AUC: 0.72) is superior or comparable to five state-of-art ICB response predictors that emphasize resistance: TIDE (average AUC: 0.70), IMPRES (average AUC: 0.62), MPS (average AUC: 0.47), Cytotoxic signature (average AUC: 0.72), and functional ICB resistance (average AUC: 0.58) (**Fig. 2C**). We next benchmarked our RDS against other established transcriptomic markers of ICB therapy and immune response, including *PD1*, *PDL1*, *CTLA4*, and T-cell exhaustion. These signatures exhibited high variability in their predictive performance while RDS consistently maintained considerable predictive power (**Sup Fig. 1B**). From a translational perspective, RDS accurately classifies responders and non-responders with a high odds ratio (average odds ratio: ∼2.6; see **Fig. 2D**). More importantly, RDS significantly stratifies pre-treatment patients’ survival outcomes with ICB therapy. **Figure 2E** shows the overall survival differences between low-risk (RDS score > median; n=134) and high-risk groups (RDS score <= median; n=140) across all ICB cohorts (log-rank p-value: 0.0095). Similarly, **Figure 2F** demonstrates significant stratification of low-risk (n=106) and high-risk groups (n=100) based on progression-free survival (log-rank p-value: 0.011). IMPRES, TIDE, and other established ICB transcriptomic signatures failed to replicate these stratifications (**Supp Fig. 2A-L**), highlighting the functional relevance of our RDIs as survival and response biomarkers. Permuting the RDI activity profile (average AUC: 0.56) or patient treatment response (average AUC: 0.49) do not replicate these performances (**Supp Fig. 1C**).

Additionally, we assessed the performance of RDS in TCGA melanoma (SKCM) by calculating an RDS score for each tumor sample based on RDIs learned in the ICB cohorts. We find that RDS can effectively stratify low-risk and high-risk patients based on both overall survival (**Fig. 2G**) and progression-free survival (**Fig. 2H**), with log-rank test p-values of 8.4e-5 and 0.039, respectively. This suggests that RDIs reflect fundamental immune response mechanisms impacting patients’ clinical outcome also in the absence of ICB therapy.

To study the role of RDIs in resistance development via single-cell transcriptomics, we analyzed a single-cell ICB cohort from Jerby-Arnon et al.’s study (11). By harnessing an in-house tool that we’ve implemented, we inferred activated ligand-receptor interactions from the single-cell transcriptomics data (see Methods). *Without any additional training,* we directly derived the RDS score for each patient using RDIs inferred from bulk ICB cohorts (Methods). The RDS scores in treatment naïve samples are significantly (Wilcoxon 1-sided test p-value: 2.2e-6) higher than those in post-treatment non-responding samples (**Fig. 2I**). Moreover, the derived RDS scores classify naïve and post-ICB resistant tumors with an AUC of 0.77 (**Fig. 2J**). These results further show that RDIs are predictive of ICB resistance at a single-cell resolution.

### Functional analysis reveals the potential role of resistant deactivated interactions in mediating CD8+ T cell infiltration to the TME

To investigate the possible association between resistant deactivated interactions and the development of ICB resistance, we conducted a functional analysis on the union of interactions inferred across each ICB cohort specific model. **Figure 3A** displays the 134 RDIs (out of 299 total inferred RDIs, see **Supp. Table 1** for complete RDI network) that are inferred in at least two ICB cohort specific models that we have studied. Notably, these 299 interactions within the RDI network (**Supp. Table 1**) originate from a diverse network of tumor-immune crosstalk involving all ten cell types in the tumor microenvironment (TME). Through cell type enrichment analysis, we observe significant enrichment of ligand-expressing cell types including malignant cells (Fisher’s 1-sided test p-value: 0.020; odds ratio: 1.47) and natural killer (NK) cells (Fisher’s 1-sided test p-value: 0.00014; odds ratio: 1.95) (**Figure 3B**). Enriched receptor-receiving cell types include macrophages (Fisher’s 1-sided test p-value: 0.019; odds ratio: 1.50), NK cells (Fisher’s 1-sided test p-value: 0.04; odds-ratio: 1.33), and plasmacytoid dendritic cells (pDC) (Fisher’s 1-sided test p-value: 0.001; odds-ratio: 2.29) (**Figure 3B**). The high involvement of malignant cells suggests that the tumor actively regulates the production of ligands to evade immune response. Previous studies investigating chemokines associated with lymphocyte rich TMEs in melanoma observed altered expression of CCL2, CCL3, CCL4, CCL21, CXCL10, CXCL11, and CXCL13, all of which are ligands expressed by the tumor within the RDI network (**Supp. Table 1**) (19,20). Notably, tumor derived expression of CXCL10 has been hypothesized to localize effector CD8+ T cells via their CXCR3 receptor to the tumor site, an interaction observed within our RDI network that exhibits anti-tumor properties and is associated with immunotherapy response (21,22). Additionally, the high presence of innate immunity pDC, macrophages, and NK cells testifies to their functional relevance in ICB resistance development. The innate immunity is additionally known to play pivotal roles in the initiation of adaptive immunity which results in downstream inflammation of the TME (23–25). Within our RDI network, a key interaction identified is between the CXCL9 ligand and the CXCR3 receptor lying on dendritic cells and CD8+ T cells respectively, which has been reported to enhance intratumorally CD8+ T cell response in the context of PD1 blockade in melanoma (26). Furthermore, we observe an NK cell derived expression of XCL1 interacting with XCR1 receptor on dendritic cells, which has been associated with anti-PD1 response in melanoma (27). Analysis of the enriched cell pairs further reveals the involvement of antigen-presenting cells, also showing that CD4+ T cells are actively interacting with NK cells and pDC cells (**Supp. Fig. 2A**). Collectively, our results suggest that the crosstalk between the tumor and the immune system may be disrupted and deactivated as a mechanism for evading immune response that leads to resistance to ICB treatment.

**Figure 3:**
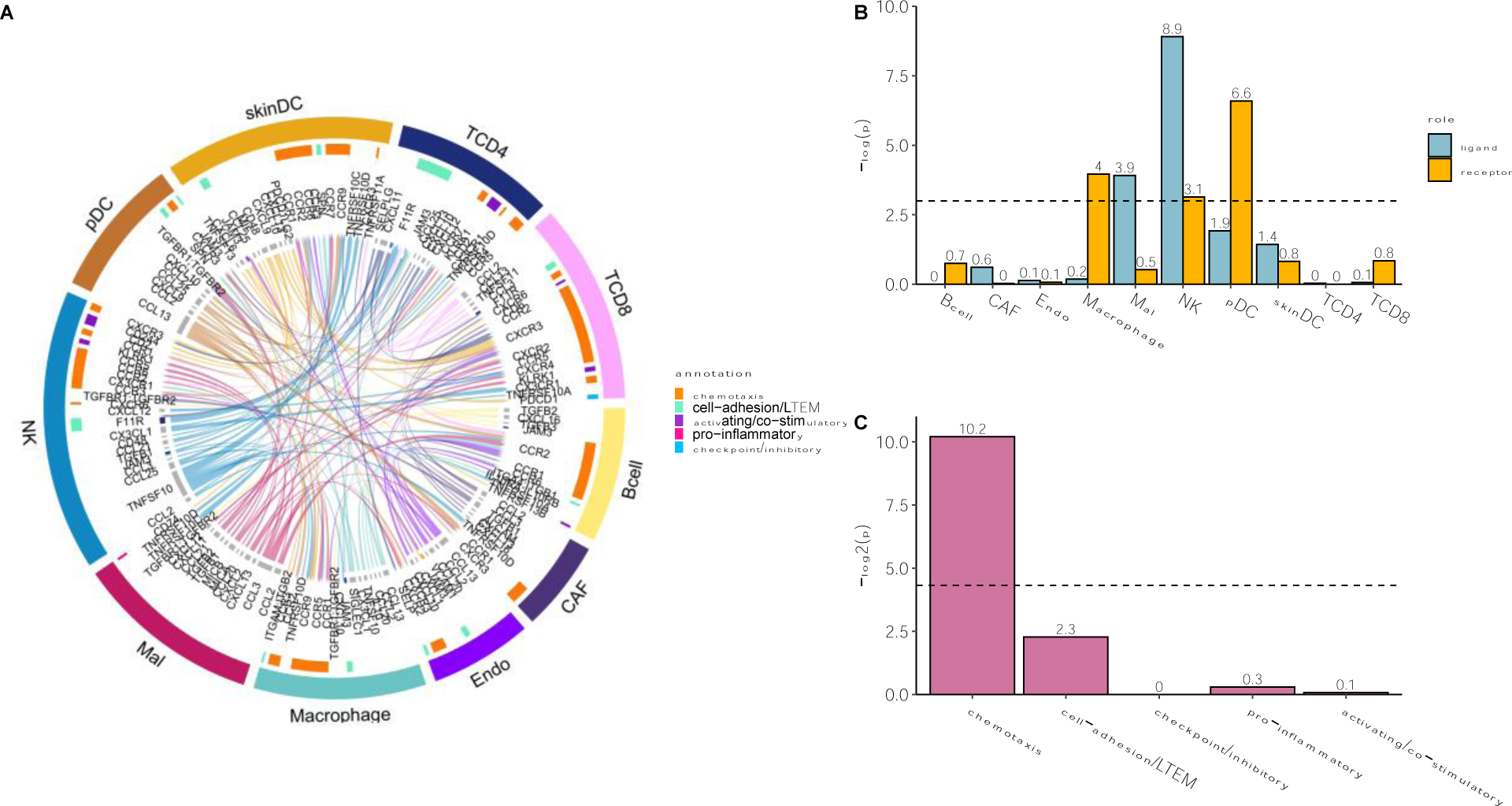
Resistant deactivated interactions are enriched in those known to mediate CD8+ T cell infiltration to the TME. **(A)** LIRICS’ chord diagram representing the 134 interactions (out of 299 interactions in the RDI network) that are overlapped in at least two ICB cohort-specific models. Cell type abbreviations detailed in the methods. **(B)** Bar plot depicting enrichment of individual cell types within RDI network as ligand-expressing or receptor-expressing cells (the background is all LIRICS’ tumor-immune interactions; N=3776). Cell types with enrichment p-values < 0.05 (dotted line) are shown. **(C)** Bar plot depicting enrichment of functional annotations within RDI network (background is the same as above). Functional annotations with p-values < 0.05 (dotted line) are shown.

Notably, reviewing the functional enrichment of the RDIs, there is a prominent over-representation of chemotaxis interactions (Fisher’s 1-sided test p-value: 0.0008; odds ratio: 3.21; **Fig. 3C**). Notably, these include interactions that have been previously shown to be associated with the recruitment and trafficking of effector CD8+ T cells to the tumor mass including CXCL9,10,11 expressed on tumors, cancer associated fibroblasts, and trafficking CD8+ T cells via their CXCR3 receptor (21,25,28); CX3CL1 expressed within the TME recruiting CD8+ T cells via their CX3CR1 receptor (26); CXCL12 expressed in tumors and involved in homing effector CD8+ T cells via their CXCR4 receptor (25,29); CXCL16 ligand expressed by cancer associated fibroblasts cells attracting effector CD8+ T cells via their CXCR6 receptor (28) and finally, CCL3,5 from NK and dendritic cells, which in turn activates CCR5 receptors on dendritic cells that recruit CD8+ T cells downstream (28).

Beyond examining the previous pertaining literature, we proceeded to further study the role of resistant deactivated interactions (RDIs) in recruiting CD8+ T cells by studying the association between the RDS scores and T-cell infiltration levels in the TME of TCGA melanoma (SKCM) samples (which has not been included in the training cohort inferring the RDS scores). Our analysis shows a strong positive correlation (Pearson correlation: 0.65; p-value < 2e-16) between RDS score and the fraction of CD8+ T cells (inferred via CODEFACS, Methods) across TCGA melanoma samples (**Fig. 4A**). Concomitantly, there is a strong negative correlation (Pearson correlation: −0.74; p-value < 2e-16) between RDS score and the fraction of malignant cells (**Fig. 4B**). We further examined the association between the RDS score and reported transcriptomic signatures of T-cell infiltration (**Fig. 4C**), T-cell exclusion (**Fig. 4D**), and post-resistance to ICB (**Fig. 4E**) in melanoma (11). Notably, we find that RDS is significantly positively correlated (Pearson correlation: 0.78; p-value < 2e-16) with the T-cell infiltration signature, while exhibiting negative correlations with the signatures of T-cell exclusion (Pearson correlation: −0.73; p-value < 2e-16) and post-ICB resistance (Pearson correlation: −0.72; p-value < 2e-16). Signatures of exclusion and resistance were all controlled for T-cell infiltration as recommended by Jerby-Arnon et al. (11) for more accurate prediction.

**Figure 4:**
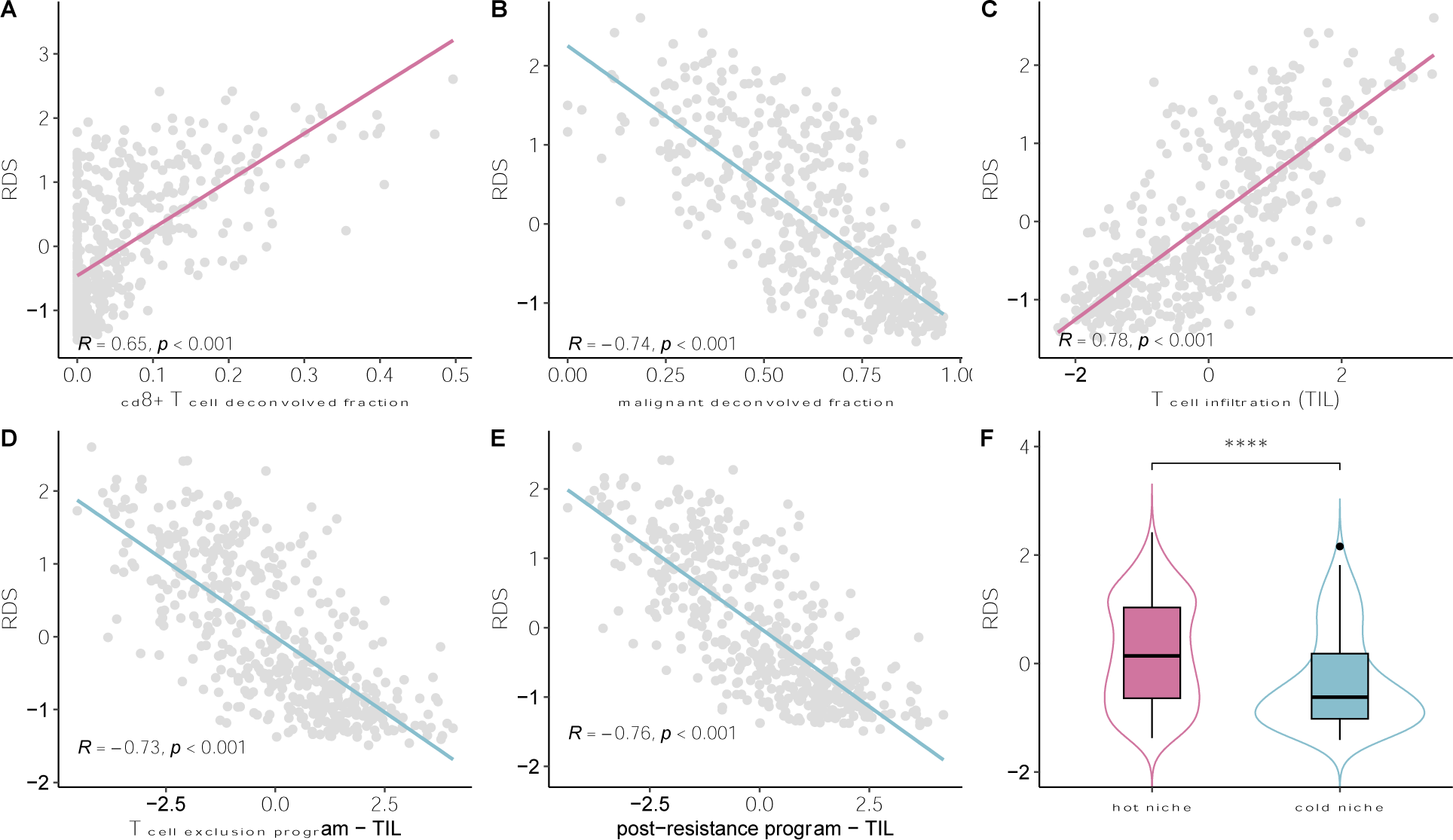
RDS scores are associated with measures of T-Cells infiltration into the TME. (A) A scatter plot depicting the correlation between RDS and estimated CD8+ T cell fraction from CODEFACS in TCGA-SKCM. (B) Scatter plot depicting the correlation between RDS and estimated tumor (malignant) fraction from CODEFACS in TCGA-SKCM. (C) Scatter plot depicting the correlation between RDS and transcriptomic signatures of T-cell infiltration (TIL) (11) in TCGA-SKCM. (D) Scatter plot depicting the correlation between RDS and transcriptomic signatures of T-cell exclusion (11) in TCGA-SKCM. The X-axis indicates the T-cell exclusion program score adjusted for TIL. (E) Scatter plot depicting the correlation between RDS and transcriptomic signatures of post-ICB resistance program (11) in TCGA-SKCM. The X-axis indicates the post-ICB resistance program adjusted for TIL. (F) Boxplot depicting distribution of RDS between hot (brisk) and cold (non-brisk) tumor niches in TCGA-SKCM.

Additionally, we studied the association between RDS and T-cell infiltration as quantified by expert pathologists’ assessment of density and arrangement of infiltration in TCGA tumor slides. Tumors with broad intra-tumoral lymphocyte infiltration were termed as “brisk” patterns testifying to immunological “hot” TMEs, while tumors with partial or more focal lymphocyte infiltration are termed as “non-brisk” patterns and “cold” TMEs (30), where hot/brisk tumor niches have been associated with favorable prognosis in melanoma (30–32) (**Figure 4E**). Based on these annotations we find that patients classified with hot tumor niches have significantly (Wilcoxon 1-sided test p-value: 8.1e-8) higher RDS scores than those with cold tumors (**Figure 4F; Supp. Figure 5E**). Without any additional training, the RDS score classifies hot vs. cold tumors with an AUC of 0.66 in TCGA-SKCM (**Supp. Figure 5F**).

Collectively, our findings suggest that RDS is predictive of ICB therapy response and the identified RDIs are those involved in trafficking CD8+ T cells to the tumor site. Initially, those identified RDIs play a crucial role in recruiting CD8+ T cell to the tumor site and maintaining an active immune infiltrating environment sustainably. Pre-treatment activation of RDIs results in high CD8+ T cell infiltration levels in the TME reinforcing a hot tumor niche (**Fig. 5A**). Following ICB treatment, checkpoint interactions are blocked, leading to the transition of CD8+ T cells into a cytotoxic state. Consequently, the tumor size is reduced (**Fig. 5B**). To evade immune attacks, resistant tumor depletes active RDIs, disrupting the machinery trafficking lymphocytes to the TME via cell-cell commutation-mediated mechanisms. This reduces the lymphocyte infiltration, converting the previously “hot” TME to “cold” (**Fig. 5C**).

**Figure 5:**
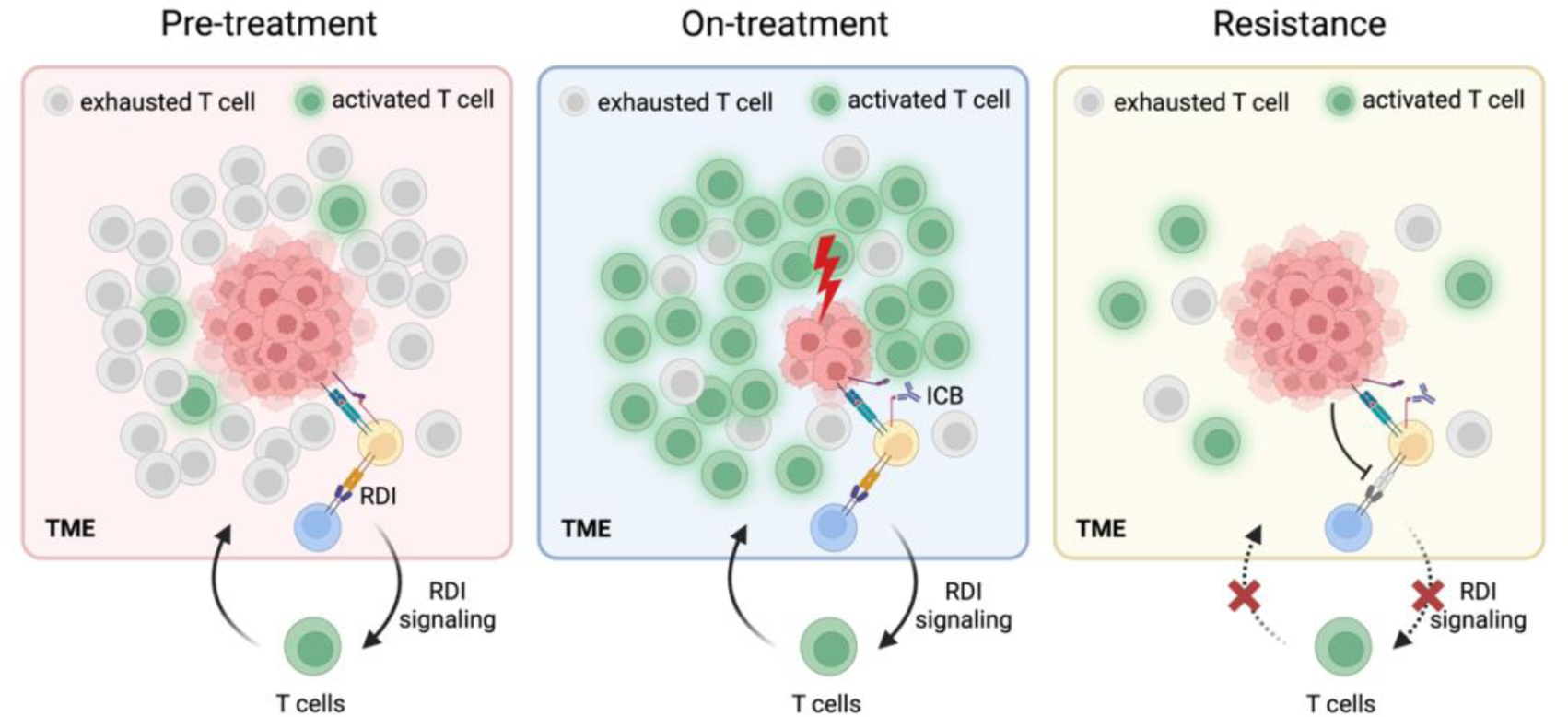
A hypothesized ICB resistance development mechanism. (**Pre-treatment**)In the pre-treatment phase, activated RDIs in the TME support the trafficking of CD8+ T cells, sustaining a hot tumor niche. However, presence of active checkpoint interactions renders the CD8+ T cells in an exhausted state, enabling tumor growth. (**On-treatment**) In the on-treatment phase, blocked checkpoint interactions empower cytotoxic CD8+ T cells within the hot tumor niche, effectively diminishing the tumor size. (**Resistance**) In resistant tumors, the tumor evades immune detection by suppressing RDIs, transitioning to a cold TME. Hindered CD8+ T cell trafficking fosters continued tumor growth, resisting checkpoint therapy.

## Discussion

Identification of resistance factors relevant to cancer immunotherapy can enhance our understanding of resistance mechanisms and uncover novel therapeutic targets. Here, we present IRIS, a new computational tool that identifies ICB resistance associated ligand-receptor interactions in the tumor microenvironment (TME). Analyzing five large melanoma cohorts, we identify such predictive interactions, that when deactivated are associated with ICB resistance. An Independent validation in a single-cell ICB cohort further supports these findings. These interactions are highly enriched in stimulatory chemokine recruitment of CD8+ T cells to the tumor mass and are associated with “hot” tumor niches.

Our findings suggest that the inhibited recruitment of T cells to the tumor site is mainly associated with the loss of stimulatory chemokine signaling. Accordingly, following ICB treatment, the tumor actively modulates the TME to deactivate interactions that promote CD8+ T cells infiltration, as specifically described in the pertaining Results section (21,22,25,26,28,29). Consequently, the TME transitions from a “hot” to a “cold” niche. These findings emphasize the importance of investigating stimulatory chemokine signaling in maintaining CD8+ T cell infiltration within the TME of ICB therapy responders (20,25,33,34). Indeed, our findings are aligned with recent reports of interventions aimed at enhancing stimulatory chemokine expression to improve anti-tumor response, which are currently under investigation in pre-clinical models (34–40).

The RDS score can serve as a predictive biomarker for ICB response, complementing existing transcriptomic biomarkers. Importantly, RDS offers several advantages over previous biomarkers: 1. It can be effectively applied to both bulk and single-cell transcriptomics data. 2. It exhibits robust predictive power across many different cohorts. 3. The identified interactions may uncover potential therapeutic targets, opening new avenues for therapeutic intervention. However, it is important to acknowledge the limitations of our study and identify areas for improvement. Firstly, like any predictive modeling approach, we identify associations and not causal mechanistic factors. One potential approach to address this is to incorporate additional functional layers that can further refine the selection of interactions that are more likely to be drivers. For example, recent studies have proposed methods to explore the function of downstream transcription factors that are regulated by ligand-receptor interactions (41–43). In our context, more causally related factors are expected to regulate specific transcription factors that play a crucial role in anti-tumor activity. Secondly, our study “*only”* considered well-defined ligand-receptor interactions from the existing literature. Furthermore, we did not incorporate spatial proximity information in our analysis. To address these limitations, future studies could employ approaches to infer de novo ligand-receptor interactions using single-cell transcriptomics data and focus on cell type pairs that exhibit spatial proximity. Thirdly, there is a lack comprehensive functional annotations of ligand-receptor interactions. This is particularly relevant for chemokine interactions, where the same ligand-receptor pair can mediate both stimulatory and inhibitory effects in immune trafficking across different target cell types (21,28,44). To address this knowledge gap, one potential solution is to computationally assess the association between each interaction and a clinical phenotype of interest across diverse cell types and contexts, which is of course a complex challenge on its own.

In summary, our study presents a new approach for identifying ICB resistance-relevant interactions using bulk deconvolved transcriptomics. Its application testifies that ICB resistance in melanoma is primarily associated with the deactivation of cell-type-specific interactions that regulate CD8+ T cell trafficking and infiltration within the tumor microenvironment.

## Methods

### Bulk RNA-seq datasets

We collected five publicly available bulk RNA-seq datasets from formalin-fixed-paraffin-embedded (FFPE) tumor tissue biopsies, accompanied by clinical information of melanoma patients undergoing immune checkpoint blockade (ICB) therapy. These datasets encompass Auslander et al. (n=37), Gide et al. (n=91), Riaz et al. (n=98), Liu et al. (n=119), and PUCH (n=55) datasets (8,15–18). The collection of datasets was frozen by Mar 2023. Each bulk RNA-seq dataset comprises a minimum of 30 samples, ensuring reliable deconvolution for ten distinct cell types. To ensure consistency for deconvolution using CODEFACS, we normalized gene expression values to transcripts per million (TPM) for each dataset, following the guidance of Wang et al. (14). Notably, all patients across these clinical studies received anti-PD1 monotherapy or a combination of anti-PD1 with anti-CTLA4 except Auslander et al. (8), which includes samples (n=6) treated exclusively with anti-CTLA4 monotherapy. For Auslander et al., Gide et al., and Riaz et al., both pre- and on-treatment samples were collected. For Gide et al., Riaz et al., and Liu et al., Transcript per million (TPM) expression data, along with CODEFACS’ deconvolved expression and cell fraction data for ten cell types were acquired from Wang et al. (14). For Auslander et al., expression data was downloaded from GEO (GSE115821), while expression and clinical data for PUCH was downloaded from (https://github.com/xmuyulab/ims_gene_signature). In the PUCH dataset, we converted the overall survival duration into days by multiplying the reported months by 30.

In addition, we obtained the deconvolved expression and cell fraction data utilizing CODEFACS for skin cutaneous melanoma patients in the The Cancer Genome Atlas (TCGA-SKCM) from Wang et al. (14). Overall survival and progression free interval (synonymous to progression free survival) for TCGA-SKCM patients was downloaded from the UCSC Xena browser (https://xenabrowser.net). The pathology classification (brisk or non-brisk) of TCGA-SKCM samples was acquired from Saltz et al. (30). Indeterminate (n=8) and “none” (n=5) pathology classifications from Saltz et al. were excluded from subsequent analyses.

### Deconvolution of bulk ICB RNA-seq datasets using CODEFACS

To deconvolve newly acquired ICB datasets, we utilized cell-type-specific signatures for ten distinct cell types sourced from Wang et al. (14). These signatures are tailored to melanoma tumors, encompassing key cell types that best represent the melanoma TME, comprising malignant cells (Mal), skin dendritic cells (skinDC), plasmacytoid dendritic cells (pDC), CD8+ T lymphocytes (TCD8), CD4+ T lymphocytes (TCD4), macrophages, natural killer cells (NK), B lymphocytes (Bcell), endothelial cells (Endo), and cancer associated fibroblasts (CAF). Leveraging the provided signature and bulk datasets, CODEFACS (14) estimated the cell-type-specific expression and cell fractions for the identical set of ten cell types across all five deconvolved melanoma ICB cohorts.

### Inferring “active” and “inactive” cell-type-specific ligand-receptor interactions using LIRICS

Deconvolved expression data from the five melanoma ICB cohorts and TCGA-SKCM were inputted into LIRICS (14), which infers cell-type-specific ligand-receptor interactions (LRIs), classified as “active” (1) or “inactive” (0) for each tumor sample within a given cohort. This procedure was conducted for pre-treatment samples and on-treatment samples separately. The scope of these inferred interactions is confined to those LRIs present within the original curated LIRICS database (n=3776).

### Validate resistant depleted interactions using single-cell RNA-seq dataset

We obtained single-cell RNA-seq data (scRNA-seq) encompassing both naïve/untreated (n=16) and post-ICB resistant (n=15) FFPE melanoma tumors from Jerby-Arnon et al. (11). Unfortunately, the post-ICB responder sample was omitted from our analyses due to its limited patient sample size (n=1). To remain consistent with the bulk datasets, we reannotated the cell types to the ten specified cell types previously mentioned. T cells that did not align with CD4+ or CD8+ subtypes were excluded. Given the variances in available cell types across each tumor sample, we initially consolidated all cells within each treatment group (naïve or post-ICB resistant). Subsequently, we down sampled 40% of cells for each cell type to create “pseudo-samples” corresponding to their respective treatment group. This procedure was repeated through 200 iterations for each treatment group, yielding 200 naïve and 200 post-ICB resistant pseudo tumor samples. To discern the “active” or “inactive” status of curated LIRICS interactions using scRNA-seq data, we adopted the method provided by Kumar et al. (45) with interactions exhibiting an empirical p-value < 0.05 categorized as “active.”

### Summary of IRIS

IRIS (Immunotherapy Resistance cell-cell Interaction Scanner) operates as a supervised machine learning algorithm, following a two-step process: 1. Differential activation analysis between pre-treatment and on-treatment non-responder group to extract a set of cell-type-specific ligand-receptor interactions, and 2. Employ a hill climbing aggregative feature selection algorithm to further optimize the selected interactions from step 1 by maximizing their classification power in distinguishing responders from non-responders in the pre-treatment phase. Together, these steps aim to infer either a set of “resistance deactivated interactions” (RDI) or “resistance activated interactions” (RAI). When multiple training cohorts are available, the training cohorts used in step 1 and step 2 are mutually exclusive and iteratively exchanged resulting in an ensemble model with multiple sets of inferred RDIs or RAIs for a given testing-cohort.

### Differential activation analysis in step 1

The first step of IRIS aims to determine interactions that are either “activated” or “deactivated” following the emergence of immune checkpoint blockade (ICB) resistance. This involves using cell-type-specific ligand-receptor interactions as features, and the objective is to identify differentially activated interactions between pre-treatment and non-responder (RECIST criteria: stable/progressive disease) on-treatment samples. This step both establishes the directionality of the interactions following ICB resistance and reduces the search space for the subsequent step 2.

Training involves utilizing the cell-type-specific ligand-receptor interactions activity profile, a direct output of LIRICS. When multiple training cohorts are available, interaction activity profiles are merged, including the union of inferred interactions within each cohort. The initial feature-space can consist of up to 3776 possible ligand-receptor interactions between ten cell types within the melanoma TME, sourced from the LIRICS-curated database. A fisher’s exact test is employed to identify interactions that are significantly activated (FDR < 0.2 per cell type pair) in either the on-treatment non-responder or the pre-treatment groups, categorizing the selected interactions as either resistance “activated” or resistance “deactivated” respectively. The output of step 1 is a ranked list (by FDR values) of interactions that are categorized as either RAIs or RDIs. These interactions compose the search space for step 2.

### Hill climbing aggregative feature selection for interactions that maximize classification of patient response to ICB in step 2

The second step of IRIS aims to identify an optimal set of interactions that are functionally relevant in dictating ICB response. This involves starting from the list of interactions (RDI or RAI) identified in the previous step as features, and the objective is to select for an optimal set of interactions that maximizes the classification power in distinguishing responders (RECIST criteria: partial/complete response) and non-responders (RECIST criteria: stable/progressive disease) in the pre-treatment phase. The method of step 2 was developed by amalgamating components of the feature selection methodologies outlined in our previous work from Wang et al. (14) and Auslander et al. (8).

The algorithm consists of an iterative procedure comprising three sequential stages. First, the pre-treatment samples in the training cohort are randomly split into three folds: two for training and one for testing. Second, starting from an empty set, the procedure greedily adds the most effective discriminating interactions (ranked by FDR by cell-pair from step 1) in a step-by-step manner until no further improvement in the classification accuracy (in terms of AUC) of ICB response is achieved within the training-folds. Considering the directional nature of individual features established in stage 1, we anticipate that for RAIs from step 1, less activations correspond to a response. Conversely, for RDIs, higher activation levels correlate with a response. Third, the selected set of interactions from the training folds *together* is evaluated in terms of classification accuracy and scored on the testing-fold. A test AUC >= 0.6 results in a reward score of 1 for all selected set of interactions, while a test AUC < 0.4 incurs a penalty score of −1. AUCs in-between and all unselected features received a score of 0. This process is repeated for 500 iterations.

Following 500 iterations, we sum the scores for each individual interaction across all iterations, termed feature score. We identify the features with the greatest feature score from the entire 500 solutions, which approximates a solution that minimizes the risk of overfitting. During the feature selection, it is possible to estimate the empirical p-value linked to each feature having a higher feature score than the feature score obtained by random chance. This estimation is accomplished by shuffling the values of each solution derived from the hill climbing feature selection, repeating this procedure 1000 times for each iteration. By performing 1000 random permutations of 500 solutions, a null distribution representing the feature score via random chance is constructed. We selected features with an empirical p-value < 0.05 as either our RDIs or RAIs depending on the input features directionality. The empirical p-value for each feature j is estimated as follow:

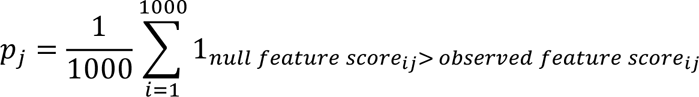

In this study, the pre-treatment cohort from Auslander et al. was excluded during step 2 feature selection, owing to the limited number of responders, which hindered the reliable determination of prediction accuracy.

### Calculating RDS and RAS

Following the identification of RDI or RAI in melanoma, the calculation of each tumor samples’ resistance-depleted score (RDS) and resistance-activated score (RAS) for a given tumor sample is computed as followed:

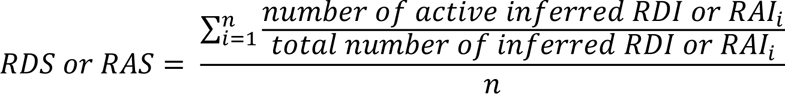

First calculate the fraction of active inferred interactions derived from each set of RDIs or RAIs inferred (i) within the ensemble model. Then average the fraction of active inferred interactions across all (n) sets of RDIs or RAIs within the ensemble model. The ultimate RDS and RAS scores are scaled for cross-cohort comparisons.

A merged ensemble model is also derived by combining all individual RDI ensemble models originally inferred for each cohort: Gide et al., Liu et al., Auslander et al., Riaz et al., and PUCH. This unified merged model is applied to the two independent patient cohorts: TCGA-SKCM and Jerby-Arnon et al.

### Calculating odds ratio

To establish the proper classification threshold for distinguishing true responder and true non-responder from false responder and non-responder for each testing cohort, we considered both the set of RDIs inferred, and tumor samples extracted from each pre-treatment training cohort used in step 2 are considered. For each training cohort, the proportion of active inferred interactions in each sample are calculated and scaled across all tumor samples. Using the scaled score in conjunction with response information for each tumor sample in the training cohort, we compute the optimal maximizing cut point using the cutpointr() function from the cutpointr package in R (46). The final threshold is obtained by taking the mean of the individual thresholds computed using each training cohort.

## Acknowledgment

This research was supported in part by the NIH Intramural Research Program, National Cancer Institute. This work utilized the computational resources of the NIH HPC Biowulf cluster (http://hpc.nih.gov). The results here are in part based upon data generated by the TCGA Research Network: https://www.cancer.gov/tcga. We would additionally like to acknowledge members of the Cancer Data Science Laboratory for their helpful feedback on this work.

Figures 1 and 5 were created with Biorender.com (biorender.com/)

## Supplementary figures

**Supplementary Figure 1:**
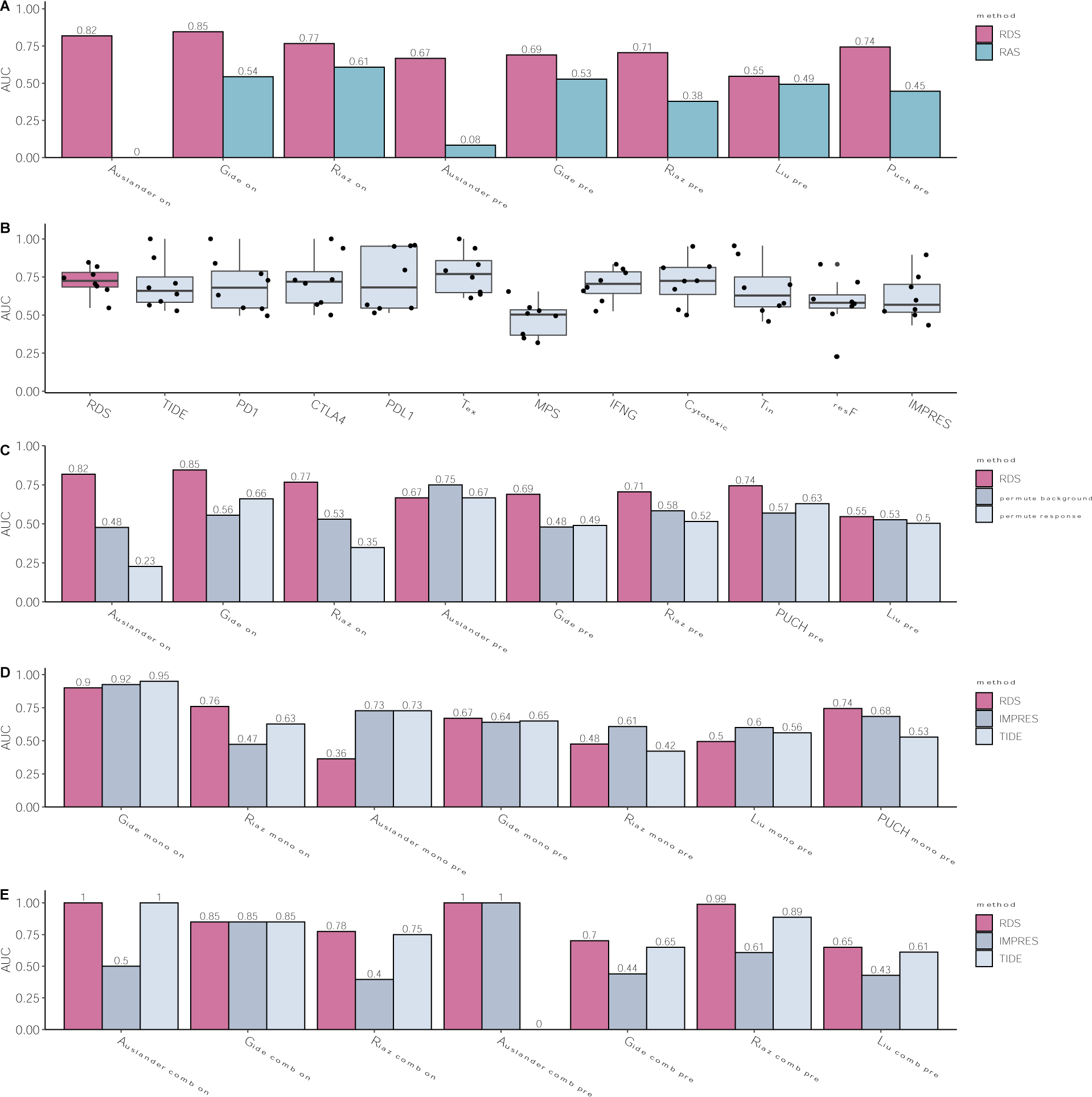
(A) Bar plot depicting AUC in classifying responder vs. non-responder melanoma samples between resistance deactivated (RDS) and resistance activated (RAS) interactions. (B) Boxplot depicting the distribution of AUC in classifying responder vs. non-responder samples in all melanoma ICB cohorts between RDS and 10 relevant transcriptomic signatures of ICB and immune response: TIDE (7), *PD1, CTLA4, PDL1*, T-cell exhaustion (Tex), Melanocytic plasticity score (MPS) (9), IFNG signature (47), Cytotoxic signatures (10), T-cell inflamed GEP (Tin) (48), functional ICB resistance (resF) (11), IMPRES (8). (C) Bar plot depicting AUC in classifying responder vs. non-responder melanoma samples between RDS, RDS when permuting cell-type-specific ligand-receptor interaction profile, and RDS when permuting patient response. (D) Bar plot depicting AUC in classifying responder vs. non-responder melanoma ICB monotherapy (anti-PD1 or anti-CTLA4) samples between RDS, TIDE (7), and IMPRES (8). (E) Bar plot depicting AUC in classifying responder vs. non-responder melanoma ICB combination therapy (anti-PD1 with anti-CTLA4) samples between RDS, TIDE (7), and IMPRES (8).

**Supplementary Figure 2:**
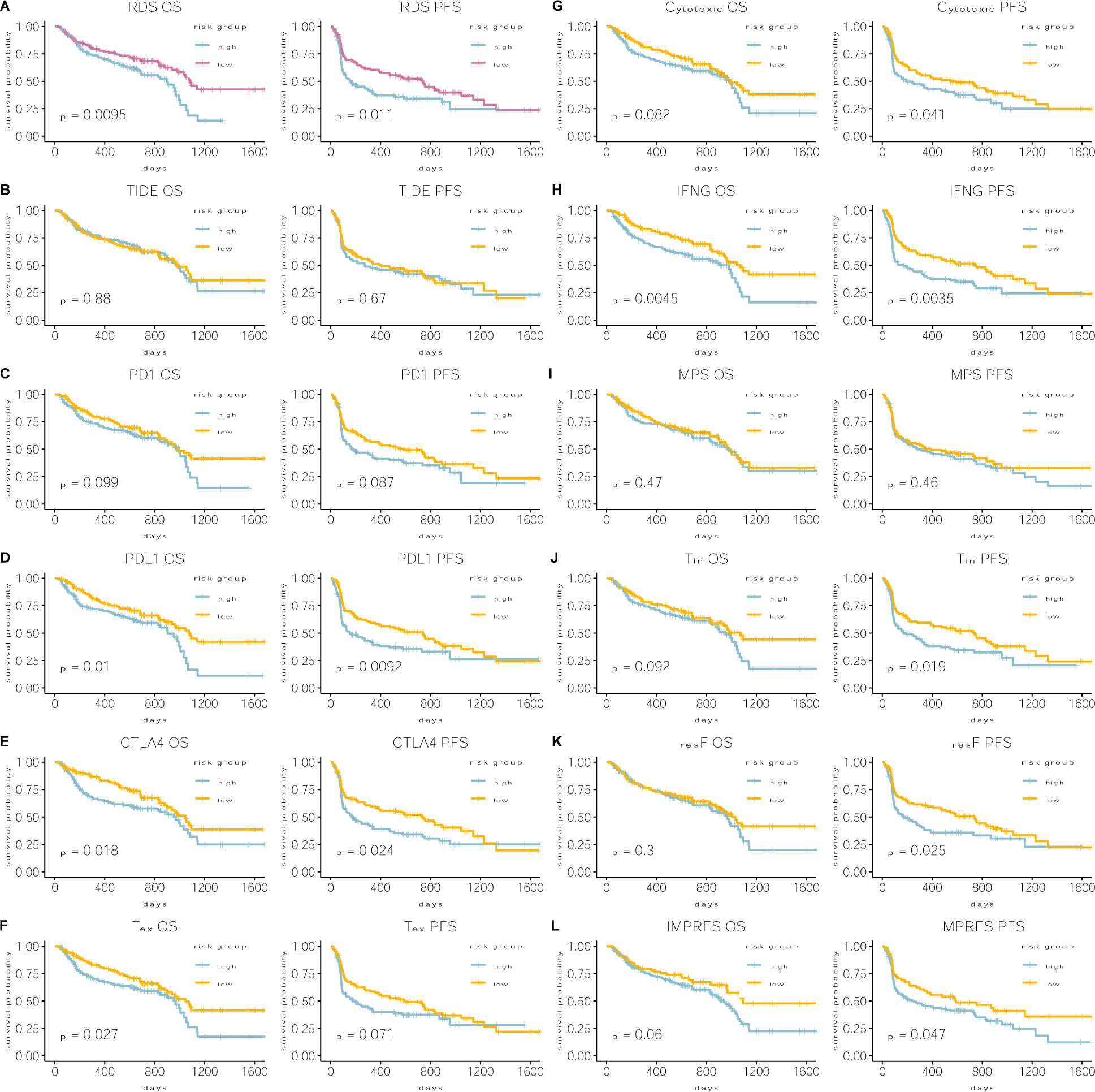
Survival stratification performance of RDS score vs. other relevant bulk-transcriptomics signatures of the combined set of pre-treatment melanoma samples receiving immune checkpoint blockade. (A) Kaplan-Meier plots depicting progression free (PFS) and overall survival (OS) differences between the low and high-risk groups defined by the median value of RDS. The significance of survival differences was estimated using the log-rank test. Time on the X-axis is measured in days. (B-L) Kaplan-Meir plots showing PFS and OS differences between low and high-risk groups defined by the median value of relevant transcriptomic signatures. The significance of survival differences was estimated using the log-rank test. Time on the X-axis is measured in days. The signatures evaluated in each panel are: (B) TIDE (7), (C) *PD1*, (D) *CTLA4*, (E) *PDL1*, (F) T-cell exhaustion (Tex), (G) Melanocytic plasticity score (MPS) (9), (H) IFNG signature (47), (I) Cytotoxic signatures (10), (J) T-cell inflamed GEP (Tin) (48), (K) functional ICB resistance (resF) (11), (L) IMPRES (8).

**Supplementary Figure 3:**
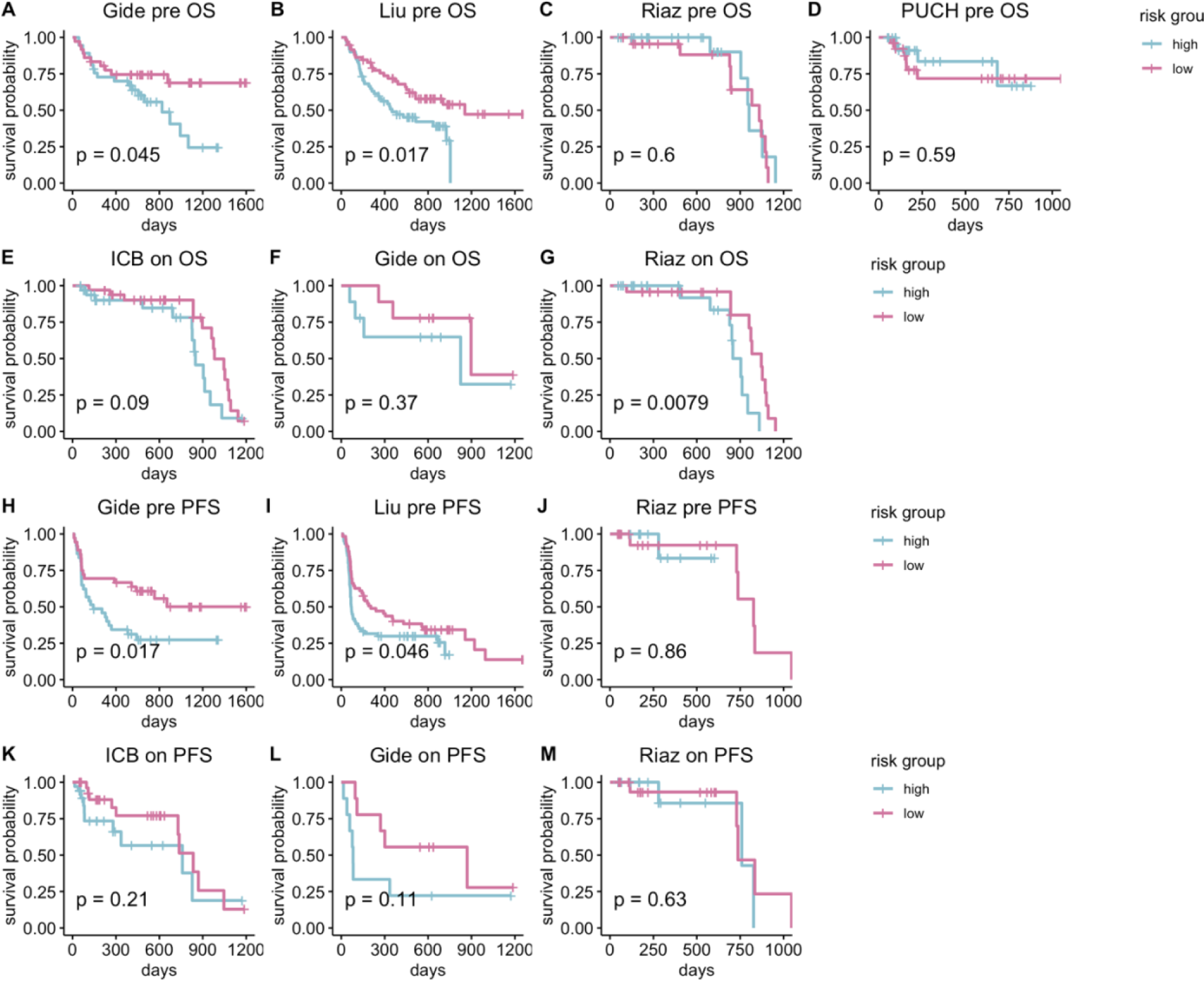
Survival stratification performance of RDS for melanoma patients receiving checkpoint therapy. Kaplan-Meir plots showing progression free survival (PFS) and overall survival (OS) differences between low and high-risk groups defined by the median value of RDS. The significance of survival differences was estimated using the log-rank test. Time on the X-axis is measured in days. (A-D) Kaplan-Meir plots depicting OS of individual checkpoint cohorts’ pre-treatment patients. (E) Kaplan-Meir plots depicting OS of the combined set on-treatment checkpoint treated patients. (F-G) Kaplan-Meir plots depicting OS of individual checkpoint cohorts’ on-treatment patients. (H-J) Kaplan-Meir plots depicting PFS of individual checkpoint cohorts’ pre-treatment patients. (K) Kaplan-Meir plots depicting PFS of the combined set on-treatment checkpoint treated patients. (L-M) Kaplan-Meir plots depicting PFS of individual checkpoint cohorts’ on-treatment patients.

**Supplementary Figure 4:**
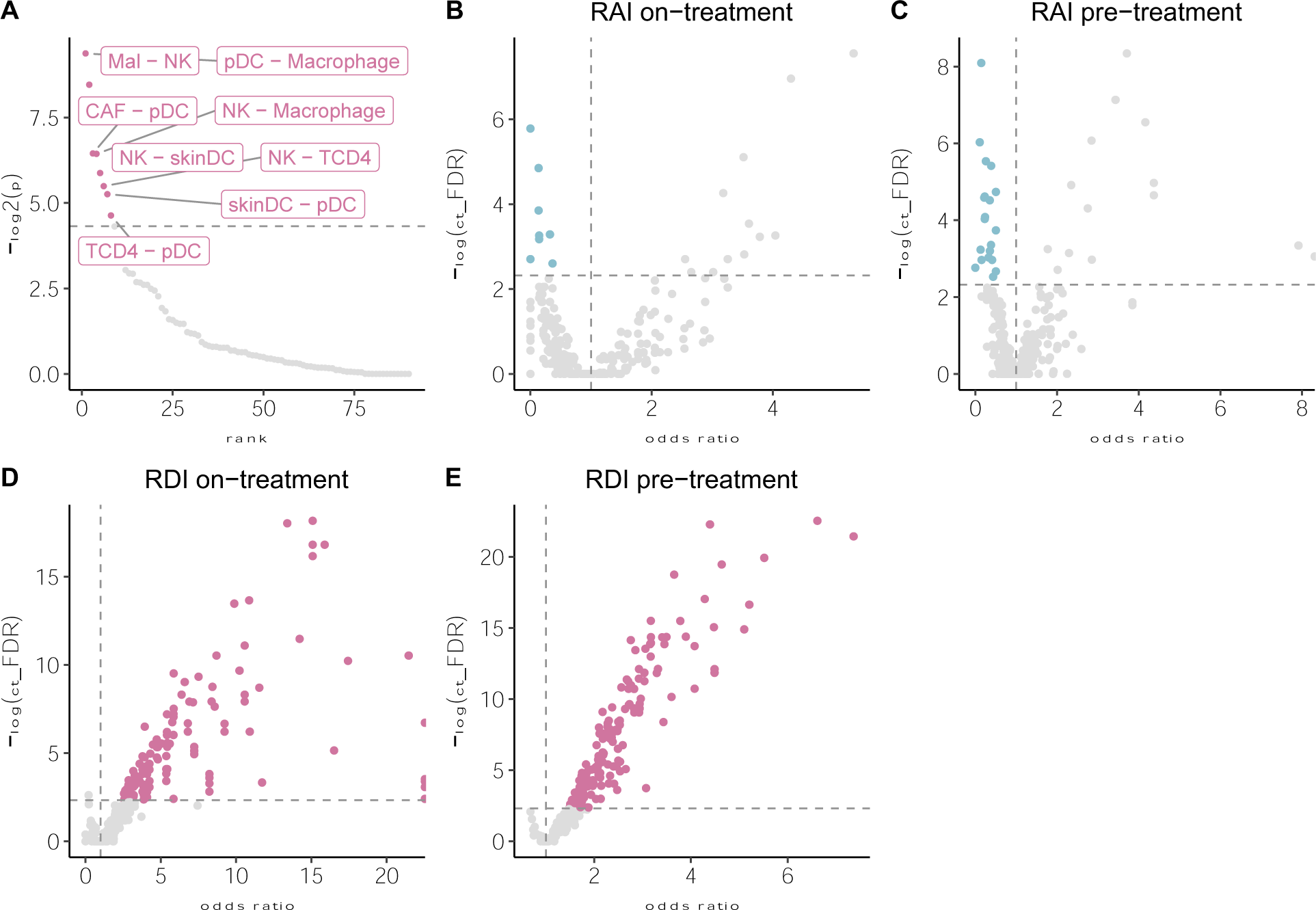
(A) Enrichment analysis depicting top ranked ligand cell-receptor cell pairs enriched within the RDI network. Background were all LIRICS’ tumor-immune interactions inferred (N=3776). The Y-axis indicates the p-value from the cell pair enrichment analysis. The X-axis indicates the rank in ascending order starting from the most significantly enriched cell-pair. Cell pairs with p-value < 0.05 (dotted line) were considered significantly enriched and are highlighted in magenta. (B) Enrichment analysis depicting individual active RAIs enrichment in responder compared to non-responder patients in the combined set of on-treatment samples receiving immune checkpoint blockade. The X-axis indicates the odds-ratio of either being enriched in the responder (> 1) compared to non-responder (< 1) samples. The Y-axis indicates the significance (FDR) of individual interactions enrichment in on-treatment responder samples (Fisher’s one-sided test). Interactions with an odds-ratio < 1 and FDR < 0.2 per cell type pair are considered significantly activated in non-responder on-treatment samples and are highlighted in blue. (C) Enrichment analysis depicting individual active RAIs enrichment in responder compared to non-responder patients in the combined set of pre-treatment samples receiving immune checkpoint blockade. The X-axis indicates the odds-ratio of either being enriched in the responder (> 1) compared to non-responder (< 1) samples. The Y-axis indicates the significance (FDR) of individual interactions enrichment in pre-treatment responder samples (Fisher’s one-sided test). Interactions with an odds-ratio < 1 and FDR < 0.2 per cell type pair are considered significantly activated in non-responder pre-treatment samples and are highlighted in blue. (D) Enrichment analysis depicting individual active RDIs enrichment in responder compared to non-responder patients in the combined set of on-treatment samples receiving immune checkpoint blockade. The X-axis indicates the odds-ratio of either being enriched in the responder (> 1) compared to non-responder (< 1) samples. The Y-axis indicates the significance (FDR) of individual interactions enrichment in on-treatment responder samples (Fisher’s one-sided test). Interactions with an odds-ratio > 1 and FDR < 0.2 per cell type pair are considered significantly activated in responder on-treatment samples and are highlighted in magenta. (E) Enrichment analysis depicting individual active RDIs enrichment in responder compared to non-responder patients in the combined set of pre-treatment samples receiving immune checkpoint blockade. The X-axis indicates the odds-ratio of either being enriched in the responder (> 1) compared to non-responder (< 1) samples. The Y-axis indicates the significance (FDR) of individual interactions enrichment in pre-treatment responder samples (Fisher’s one-sided test). Interactions with an odds-ratio > 1 and FDR < 0.2 per cell type pair are considered significantly activated in responder pre-treatment samples and are highlighted in magenta.

**Supplementary Figure 5:**
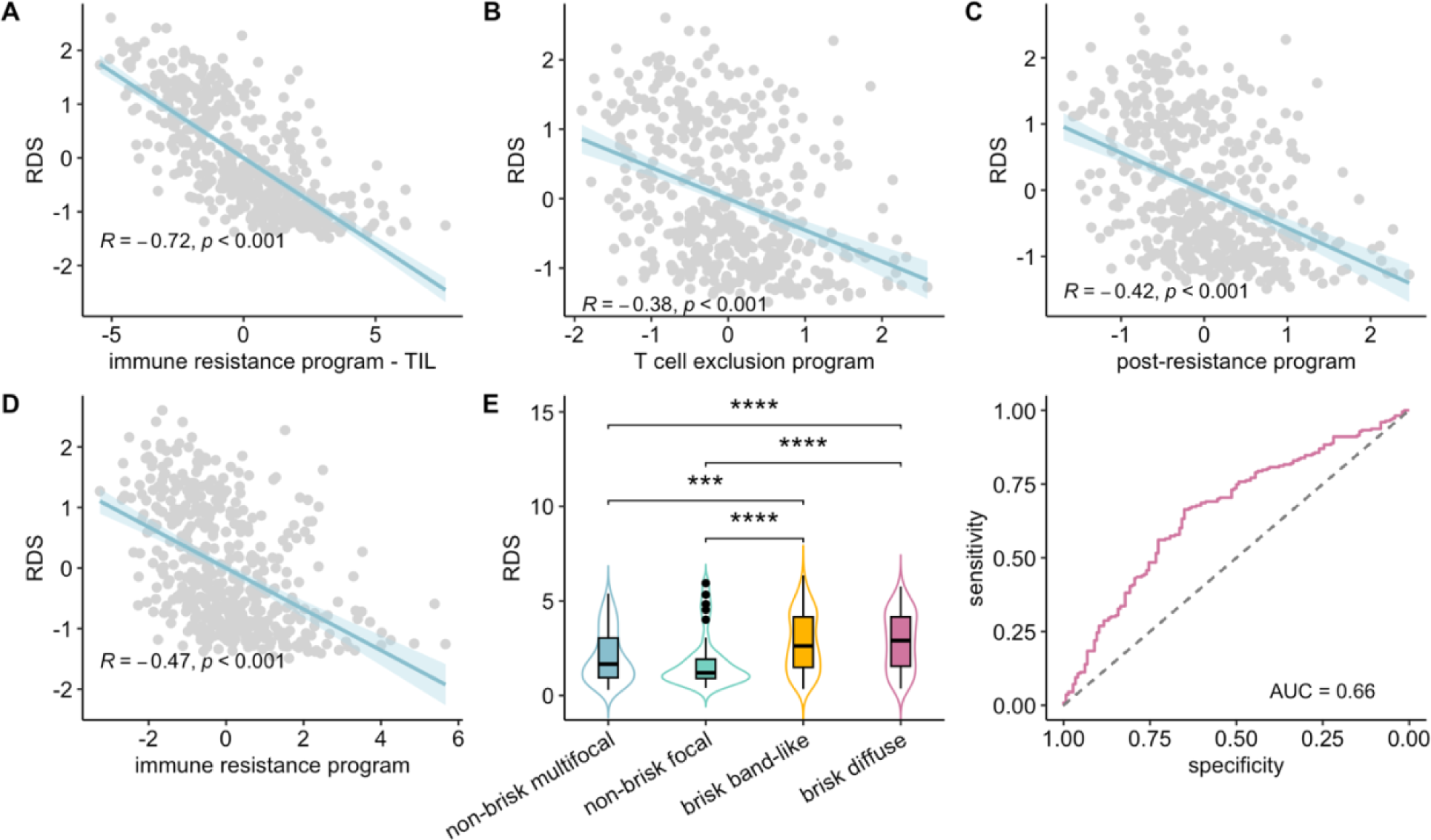
(A) Scatter plot depicting the correlation the between RDS and transcriptomic signatures of immune resistance in TCGA-SKCM. The immune resistance program includes the combined set of genes inferred for both the T-cell exclusion and post-resistance programs. The X-axis indicates the immune resistance program adjusted for TIL. The pearson correlation coefficient is 0.72 (p-value < 0.001). (B) Scatter plot depicting the correlation the between RDS and transcriptomic signatures of T-cell exclusion program in TCGA-SKCM. The pearson correlation coefficient is −0.38 (p-value < 0.001). (C) Scatter plot depicting the correlation the between RDS and transcriptomic signatures of post-resistance program in TCGA-SKCM. The pearson correlation coefficient is −0.42 (p-value < 0.001). (D) Scatter plot depicting the correlation the between RDS and transcriptomic signatures of immune resistance program in TCGA-SKCM. The immune resistance program includes the combined set of genes inferred for both the T-cell exclusion and post-resistance programs. The pearson correlation coefficient is −0.47 (p-value < 0.001). (E) Boxplot depicting distribution of RDS between non-brisk and brisk subtypes in TCGA-SKCM. (F) ROC curve depicting classifying hot vs. cold tumor niches in TCGA-SKCM using RDS.

